# Evolutionary branching of function-valued traits under constraints

**DOI:** 10.1101/841064

**Authors:** Hiroshi C. Ito

**Affiliations:** Department of Evolutionary Studies of Biosystems, School of Advanced Sciences, The Graduate University for Advanced Studies, SOKENDAI, Shonan Village, Hayama, Kanagawa 240-0193, Japan

## Abstract

Some evolutionary traits are described by scalars and vectors, while others are described by continuous functions on spaces (e.g., shapes of organisms, resource allocation strategies between growth and reproduction along time, and effort allocation strategies for continuous resource distributions along resource property axes). The latter are called function-valued traits. This study develops conditions for candidate evolutionary branching points, referred to as CBP conditions, for function-valued traits under simple equality constraints, in the framework of adaptive dynamics theory (i.e., asexual reproduction and rare mutation are assumed). CBP conditions are composed of conditions for evolutionary singularity, strong convergence stability, and evolutionary instability. The CBP conditions for function-valued traits are derived by transforming the CBP conditions for vector traits into those for infinite-dimensional vector traits.

## 1 Introduction

Biological communities are thought to have been evolving in multiple traits simultaneously. Some evolutionary traits are described with scalars or vectors, while others are described with continuous functions on spaces (e.g., shapes of organisms, resource allocation strategies between growth and reproduction along time, and effort allocation strategies for continuous resource distributions along resource property axes).

In adaptive dynamics theory, which is an extension of ESS theory for continuous strategies (Metz et al., 1996), directional evolution of function-valued traits as well as scalar and vector traits are described with an ordinary differential equation (Dieckmann and Law, 1996). Diversifying evolution of scalar and vector traits can be analyzed by examining evolutionary branching conditions (i.e., conditions for evolutionary singularity, strong convergence stability, and evolutionary instability) (Metz et al., 1996; Geritz et al., 1997, 1998; Ito and Dieckmann, 2012, 2014; Ito and Sasaki, 2016). As for function-valued traits, appropriate description of their evolution often requires constraints. Even under such constraints, directional evolution of function-valued traits can be described in an analogous manner with the case of scalar and vector traits (Dieckmann et al., 2006; Parvinen et al., 2006, 2013; Metz et al., 2016). On the other hand, as for evolutionary branching conditions, while conditions for evolutionary singularity and evolutionary stability under constraints can be derived by the calculus of variations (e.g., Pontryagin’s maximum principle) (Dieckmann et al., 2006; Parvinen et al., 2006, 2013; Metz et al., 2016), analytical treatment of convergence stability under constraints is not easy.

This paper develops conditions for strong convergence stability (Leimar, 2009) as well as evolutionary singularity and evolutionary stability, for function-valued traits under simple equality constraints, by extending the Lagrange multiplier method for vector traits (Ito and Sasaki, 2016) for infinite-dimensional vector traits. The set of conditions for evolutionary singularity, strong convergence stability, and evolutionary instability are called the candidate branching point conditions, or the CBP conditions. The next section, Section 2, assumes a single constraint and applies the Lagrange multiplier method by Ito and Sasaki (2016) for discretized function-valued traits, and extends the discretized CBP conditions for function-valued traits. Section 3 derives the CBP conditions for function-valued traits under multiple constraints. Section 4 applies the derived CBP conditions for a specific resource competition model, where individual resource utilization distribution is the function-valued trait to be analyzed. Section 5 discusses our obtained results in connection with relevant studies.

## 2 Conditions for evolutionary branching under single constraint

In this section, we consider a single constraint. Throughout the paper, italic, bold small, and bold capital letters respectively denote scalars, column vectors, and matrices.

### 2.1 Basic assumptions

To analyze evolutionary dynamics, we use adaptive dynamics theory (Metz et al., 1996). This theory typically assumes asexual reproduction, sufficiently rare mutation, and sufficiently large population size, so that the population is monomorphic and it is at an equilibrium density when a mutant emerges. When we consider a one-dimensional trait space, *x*, whether a mutant *x*′ can invade the resident population of phenotype *x*° is determined by the initial per capita growth rate of the mutant, called the invasion fitness, *f* (*x*′, *x*°). The mutant can invade the resident only when *f* (*x*′, *x*°) is positive, resulting in substitution of the resident in many cases. Repetition of such a substitution is called a trait substitution sequence, forming directional evolution toward the higher fitness. Occasionally, a mutant coexists with its resident, which may bring about evolutionary diversification into two distinct morphs, called evolutionary branching (Metz et al., 1996; Geritz et al., 1997, 1998).

The above framework can describe adaptive evolution of function-valued traits as well (Dieckmann et al., 2006; Parvinen et al., 2006, 2013; Metz et al., 2016). In this paper, we consider a function-valued trait as a distribution *s*(*z*) along a one-dimensional space *z* (its multi-dimensional extension seems easy). To denote *s*(*z*) not as its value at *z* but as a function, we introduce 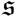. Then, a mutant and a resident are denoted by 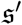 (corresponding to *s*′(*z*)) and 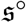 (corresponding to *s*°(*z*)), respectively. We denote the invasion fitness function by 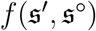. We also assume a constraint 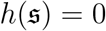, which restricts all possible fertile mutants to ones that satisfy 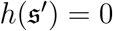.

### 2.2 Discretized function-valued traits

To analyze adaptive evolution of 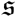 by using the Lagrange multiplier method for vector traits (Ito and Sasaki, 2016), we discretize *z* in a constant region of length *Z* into *K* sampling points *z*_0_, *z*_0_ + Δ*z*, …, *z*_0_ + *k*Δ*z*, …, *z*_0_ + [*K*− 1]Δ*z* with Δ*z* = *Z/*(*K*− 1) with a sufficiently large *K*, and express *s*(*z*) with a *K*-dimensional vector s = (*s*_1_, …, *s*_*k*_, …, *s*_*K*_)^*T*^ = (*s*(*z*_0_), …, *s*(*z*_0_ + *k*Δ*z*), …, *s*(*z*_0_ +[*K*−1]Δ*z*))^*T*^. Then we denote the invasion fitness of a mutant s′ against resident s by *F* (s′, s). The constraint is denoted by *H*(s) = 0. Note that all possible fertile mutants are restricted to the (*K*−1)-dimensional constraint surface. We see trivially that 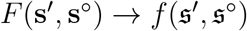 and 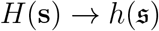 when Δ*z* → 0.

#### 2.2.1 Directional evolution

Following Ito and Sasaki (2016), we first define a Lagrange fitness function,

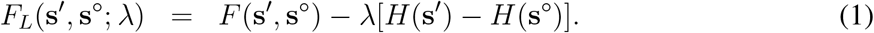

For a point s satisfying *H*(s) = 0, the λ is given by

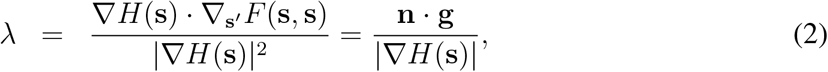

where n = (*n*_1_, …, *n*_*M*_)^*T*^ = ∇*H*(s)/|∇*H*(s)| is the normal vector of the constraint surface at s, defined by

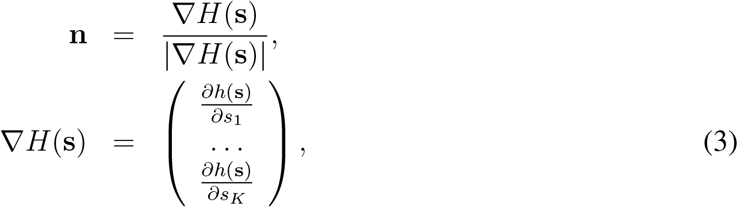

and g is the fitness gradient at s, defined by

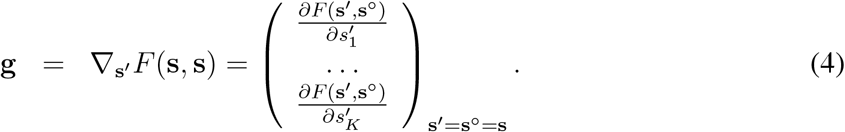

Note that the gradient of *F*_*L*_(s′, s; λ) for s is expressed as

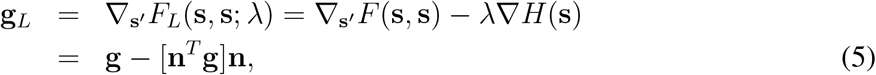

which means that g_*L*_ gives the element of g tangential to the constraint surface. When g_*L*_ = 0, there exists no fitness gradient along the constraint surface, such an s is called an evolutionarily singular point. At an evolutionarily singular point, the conditions for its convergence stability and evolutionary stability are given by the second derivatives of *F*_*L*_(s′, s°; λ) along the tangential direction of the constraint surface. Specifically, we have the following CBP conditions (Ito and Sasaki, 2016).

##### CBP conditions for discretized function-valued traits (*K*-dim traits with 1-dim constraint)

In an *K*-dimensional trait space s = (*s*_1_, …, *s*_*K*_)^*T*^, a point s is a candidate for an evolutionary branching point (CBP) along the constraint surface *H*(s) = 0, when s satisfies the following three conditions about the Lagrange fitness function, given by Eq. (1) with Eq. (2).

i. s is evolutionarily singular along the constraint surface, satisfying

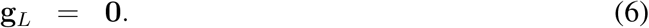
ii. s is strongly convergence stable along the constraint curve, i.e.

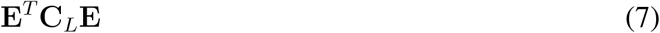

has a negative definite symmetric part.
iii. s is evolutionarily unstable along the constraint curve, i.e., a symmetric matrix

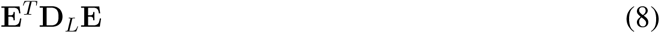

has at least one positive eigenvalue.

Here, the matrix E = (e_2_, …, e_*K*_) is composed of orthogonal base vectors of the tangent plane of the constraint surface at s, given by the diagonalization of nn^*T*^ as

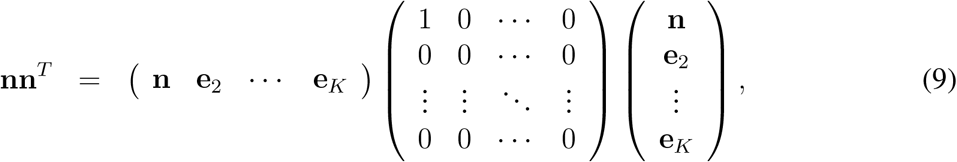

or given by the Gram Schmidt orthogonalizaiton, and

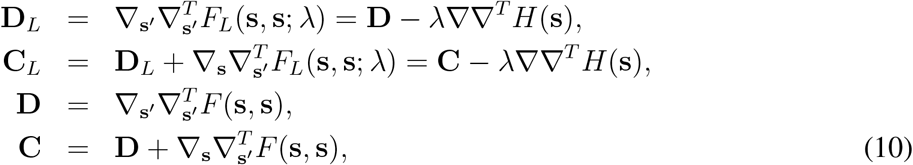

with

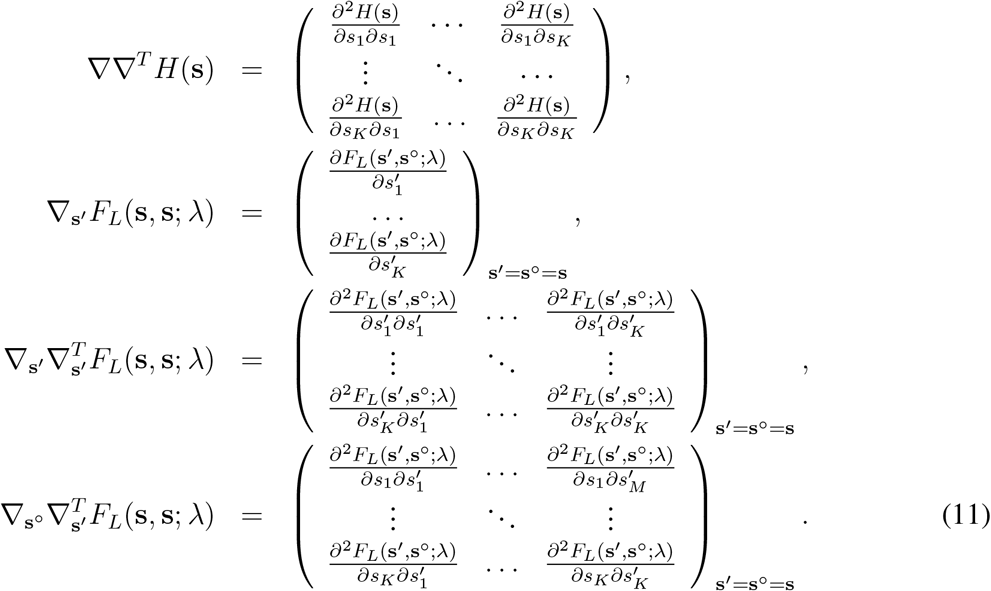

The condition (ii) and (iii) can be expressed alternatively as

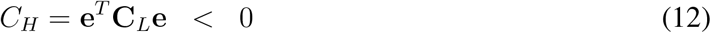

for any vector e with e^*T*^ n = 0, and

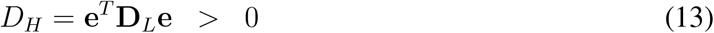

for some e with e^*T*^ n = 0, respectively.

### 2.3 Function-valued traits

Assuming Δ*z* → 0 in the above CBP conditions for discretized function-valued traits immediately gives the CBP conditions for continuous function-valued traits. Specifically, we define the Lagrange fitness function by

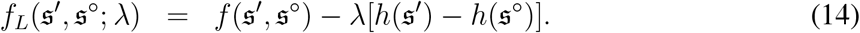

For a point 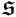, satisfying 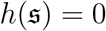 the λ is given by

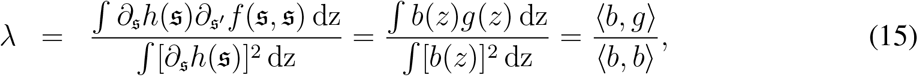

where ⟨*α, β*⟩ is the inner product operator between two distributions *α*(*z*) and *β*(*z*), defined by

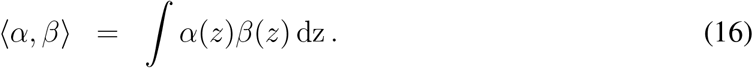

The *b*(*z*) gives the normal vector of the constraint surface at 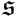, defined by

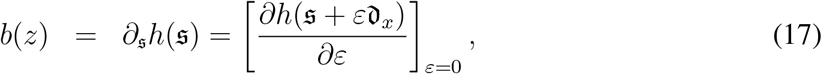

where 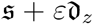 corresponds to a distribution 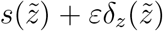 along 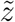, where 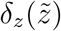 is the Dirac delta function peaked at 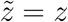. The *g*(*z*) gives the fitness gradient at 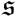, defined by

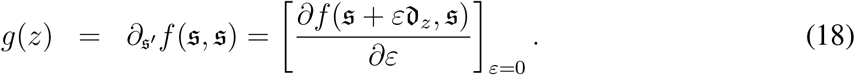

The fitness gradient at 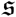 along the constraint surface is given by

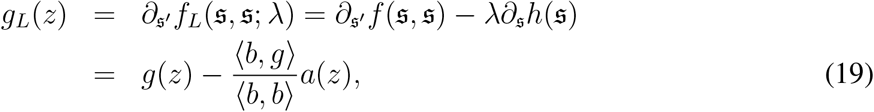

because *g*_*L*_ has no element along the constraint surface, i.e.,

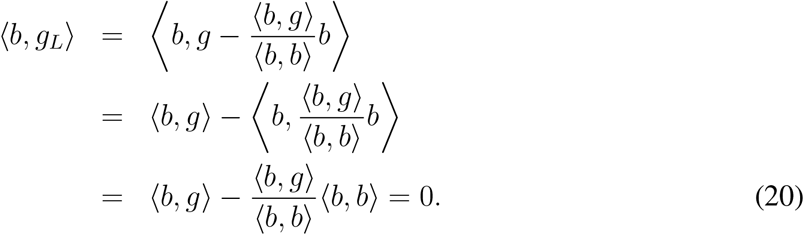

Thus, the fitness gradient for an arbitrary mutation expressed as *s*(*z*) + *εe*(*z*) without constraint is given by ⟨*e, g*⟩, while the fitness gradient for the mutation under the constraint is given by

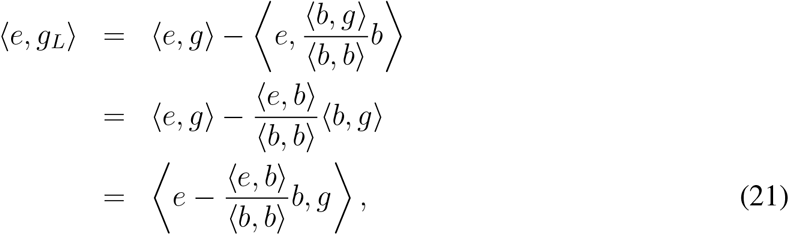

which is equivalent to ⟨*e, g*⟩ with *e*(*z*) a tangent vector of the constraint surface (i.e., ⟨*e, b*⟩ = 0).

On this basis, the CBP conditions are described as follows.

#### Theorem 1: Candidate branching point (CBP) conditions for function-valued traits (with 1-dim constraint)

In a function-valued trait space 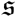, corresponding to a distribution *s*(*z*), a point 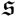 is a candidate for an evolutionary branching point (CBP) along the constraint surface 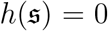, when 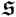 satisfies the following three conditions about the Lagrange fitness function, Eq. (14) with Eq. (15).

i. 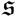 is evolutionarily singular along the constraint surface, satisfying for all *z*

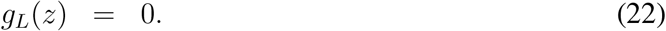
ii. 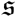 is strongly convergence stable along the constraint surface, satisfying for any *e*(*z*) with ⟨*e, b*⟩ = 0

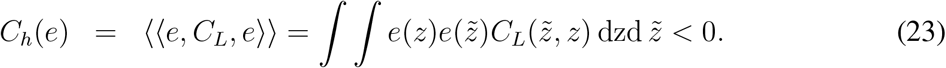
iii. 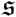 is evolutionarily unstable along the constraint surface, satisfying for some *e*(*z*) with ⟨*e, b*⟩ = 0

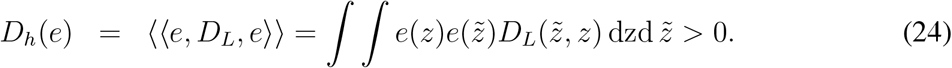

Here,

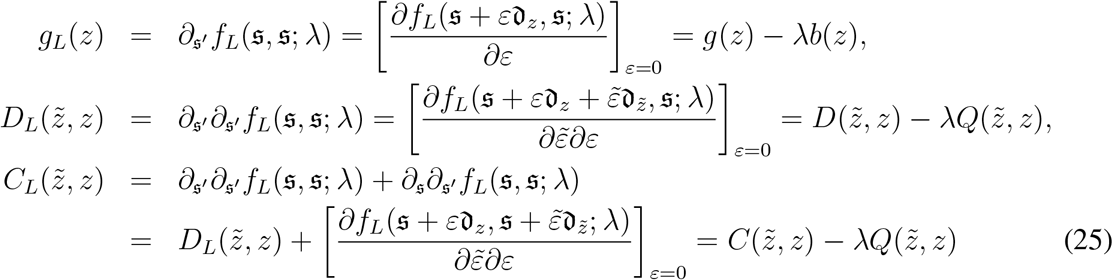

and

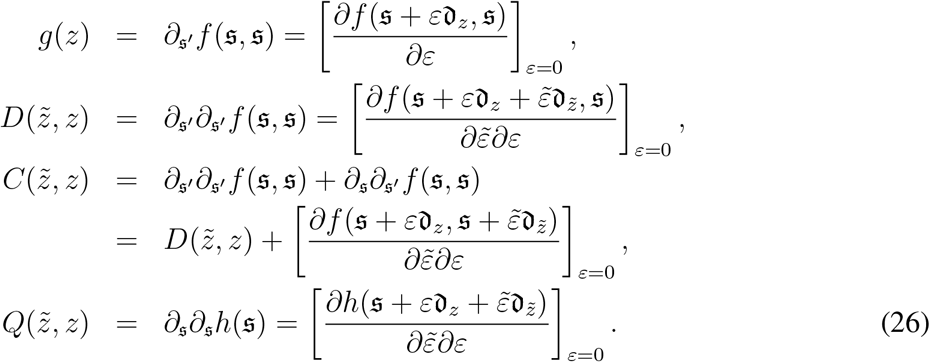

## 3 Conditions for evolutionary branching under multiple constraints

We consider multiple constraints, dented by 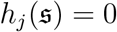 for *j* = 1, …, *N* with an arbitrary finite *N*.

### 3.1 Discretized function-valued traits

In the same manner with the previous section, we discretize *z* in a constant region of length *Z* into *K* sampling points *z*_0_, *z*_0_ + Δ*z*, …, *z*_0_ + *k*Δ*z*, …, *z*_0_ + [*K*− 1]Δ*z* with Δ*z* = *Z*/[*K* − 1] with a sufficiently large *K*, and express *s*(*z*) with a *K*-dimensional vector s = (*s*_1_, …, *s*_*k*_, …, *s*_*K*_)^*T*^ = (*s*(*z*_0_), …, *s*(*z*_0_ + *k*Δ*z*), …, *s*(*z*_0_ + *K*Δ*z*))^*T*^. Then we denote the invasion fitness of a mutant s′ against resident s by *F* (s′, s). The constraints are denoted by *H*_*j*_(s) = 0 for *j* = 1, …, *N*. Note that all possible fertile mutants are restricted to the (*K* − *N*)-dimensional constraint surface. We see trivially that 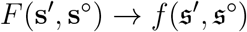 and 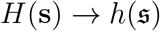 when Δ*z* → 0.

#### 3.1.1 Directional evolution

Following Ito and Sasaki (2016), we first define a Lagrange fitness function,

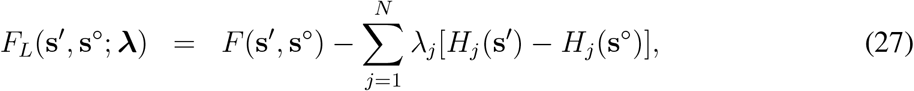

with **λ** = (λ_1_ · · · λ_*N*_). For a point s satisfying *H*_*j*_(s) = 0 for all *j* = 1, …, *N*, the **λ** = (λ_1_ · · · λ_*N*_) is given by

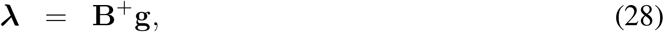

where g is given by Eq. (4), B^+^ = [B^*T*^B]^−1^B^*T*^ is the pseudo inverse of B, where

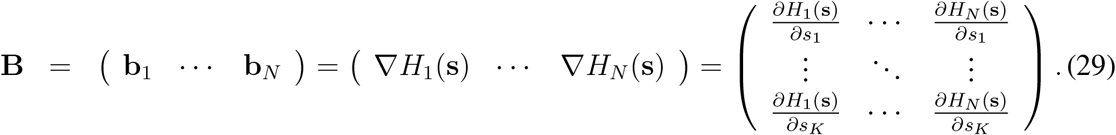

To facilitate the infinite-dimensional extension, we transform Eq. (28) as

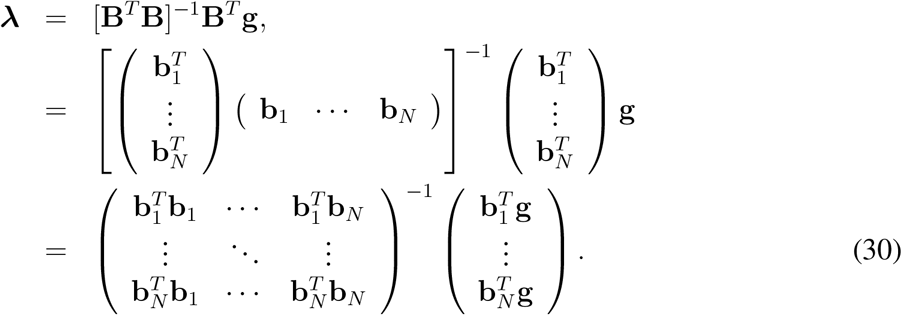

The fitness gradient at s along the constraint surface is given by

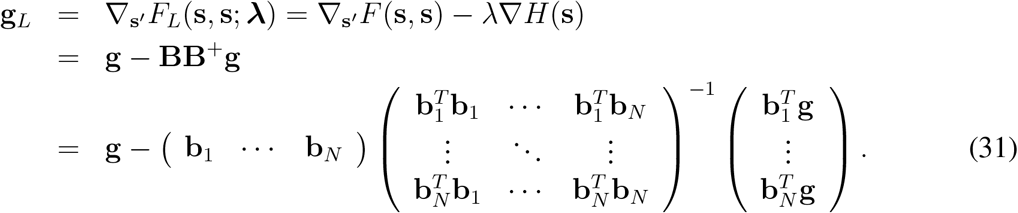

On this basis, we have the following conditions for candidate branching points.

##### CBP conditions for discretized function-valued traits (*K*-dim traits with *N*-dim constraints)

In an *K*-dimensional trait space s = (*s*_1_, …, *s*_*K*_)^*T*^, a point s is a candidate for an evolutionary branching point (CBP) along the constraint surface *H*_*j*_(s) = 0 for *j* = 1, …, *N* with *N* < *K*, when s satisfies the following three conditions about the Lagrange fitness function, given by Eq. (27) with Eq. (28).

i. s is evolutionarily singular along the constraint surface, satisfying

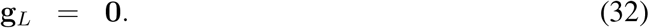
ii. s is strongly convergence stable along the constraint curve, i.e.

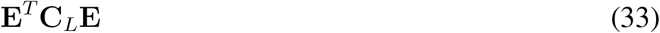

has a negative definite symmetric part.
iii. s is evolutionarily unstable along the constraint curve, i.e., a symmetric matrix

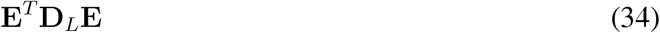

has at least one positive eigen value.

Here, the matrix E = (e_*N*+1_, …, e_*K*_) is composed of orthogonal base vectors of the tangent plane of the constraint surface at s, given by the diagonalization of BB^+^ as

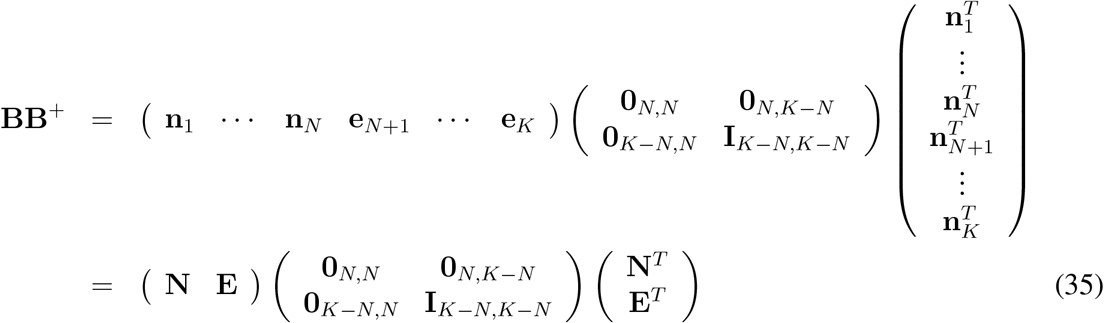

(0_*i,j*_ denotes an *i*-by-*j* zero matrix and I_*i,i*_ denotes an *i*-by-*i* identity matrix), or by the Gram Schmidt orthogonalizaiton, and

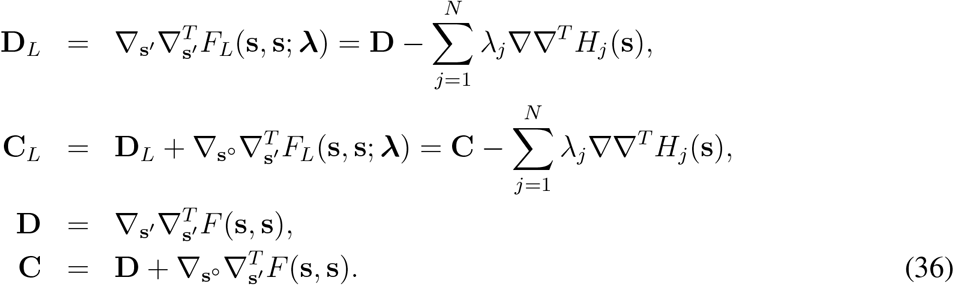

The condition (ii) and (iii) can be expressed alternatively as

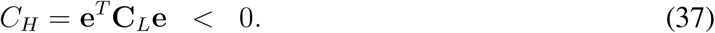

for any vector e with e^*T*^B = 0, and

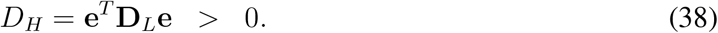

for some e with e^*T*^B = 0, respectively.

### 3.2 Function-valued traits

Assuming Δ*z* → 0 in the above CBP conditions for discretized function-valued traits immediately gives the CBP conditions for continuous function-valued traits. Specifically, we define the Lagrange fitness function by

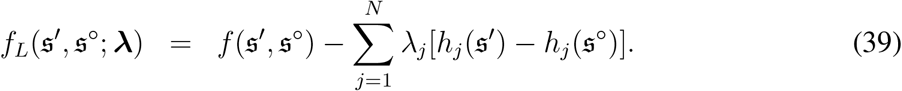

with λ = (λ_1_ … λ_*N*_). For a point 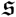 satisfying 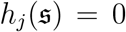 for all *j* = 1, …, *N*, the λ = (λ_1_ … λ_*N*_ is given by

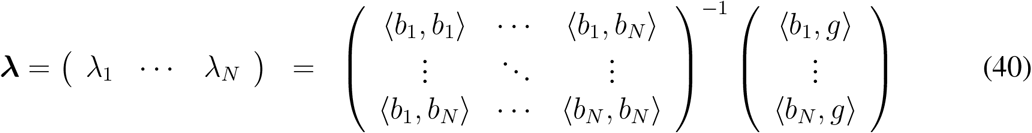

(from Eq. (30)), where

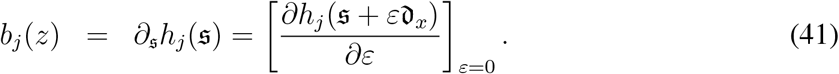

The fitness gradient at 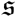 along the constraint is expressed as

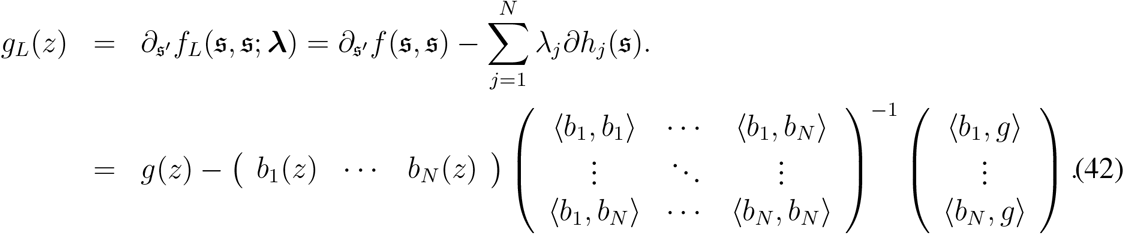

Thus, 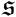 is evolutionary singular along the constraint surface when *g*_*L*_(*z*) = 0, or equivalently when ⟨*e, g*⟩ = 0 for any *e*(*z*) satisfying ⟨*e, b_j_*⟩ = 0 for all *j* = 1, …, *N*.

On this basis, we have the following conditions for candidate branching points.

#### Theorem 2: Candidate branching point (CBP) conditions for function-valued traits (with arbitrary *N* constraints)

In a function-valued trait space 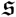, corresponding to a distribution *s*(*z*), a point 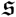 is a candidate for an evolutionary branching point (CBP) along the constraint surface 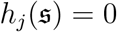 for *j* = 1, …, *N*, when 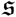 satisfies the following three conditions about the Lagrange fitness function, Eq. (39) with Eq. (40).

i. 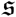 is evolutionarily singular along the constraint surface, satisfying for all *z*

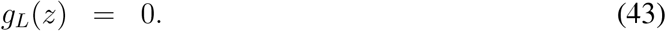
ii. 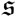 is strongly convergence stable along the constraint surface, satisfying for any *e*(*z*) with ⟨*e, b*_*j*_⟩ = 0 for all *j* = 1, …, *N*

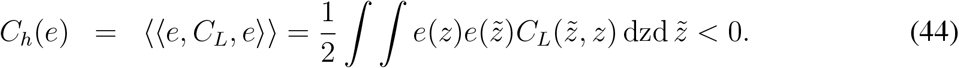
iii. 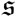 is evolutionarily unstable along the constraint surface satisfying for some *e*(*z*) with ⟨*e, b*_*j*_⟩ = 0 for all *j* = 1, …, *N*

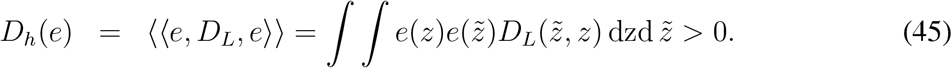

Here,

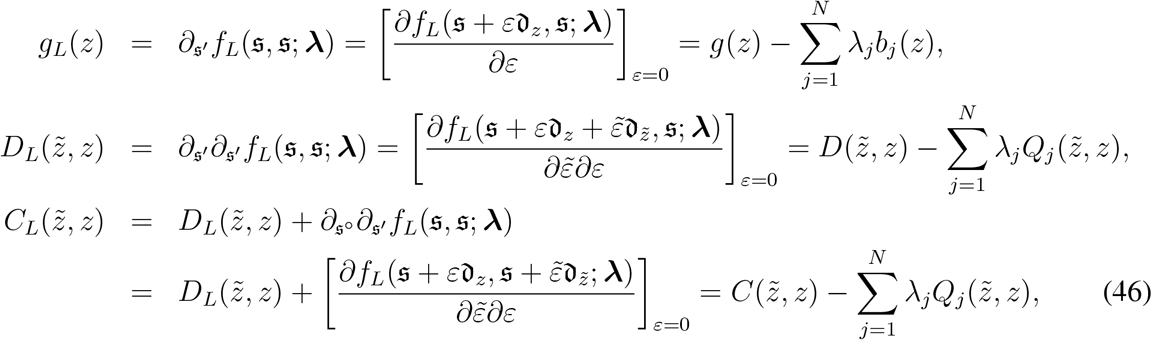

and for *j* = 1, …, *N*

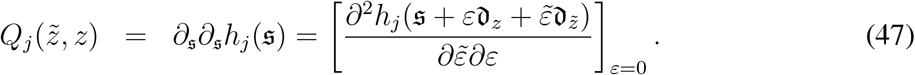

## 4 Example

We consider a resource competition model, where the distribution of individual resource consumption effort (i.e., resource utilization distribution) is the evolutionary trait, denoted by *s*(*z*) and by 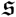. We define the dynamics of *i*th phenotype’s population density *n_i_* (under existence of *J* phenotypes) by

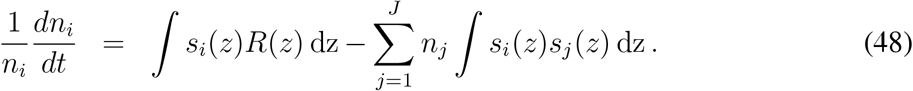

The invasion fitness of a mutant 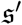 against a monomorphic resident 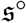 at an equilibrium density

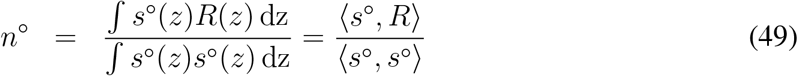

is given by

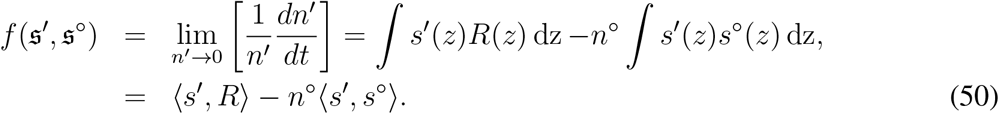

### 4.1 Evolution without constraints

We see for a point 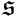,

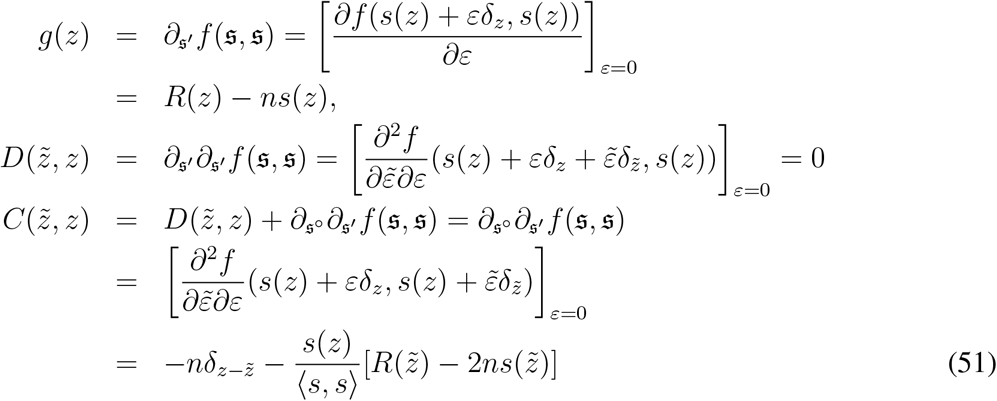

Thus, for an arbitrary mutation denoted by *e*(*z*), we get

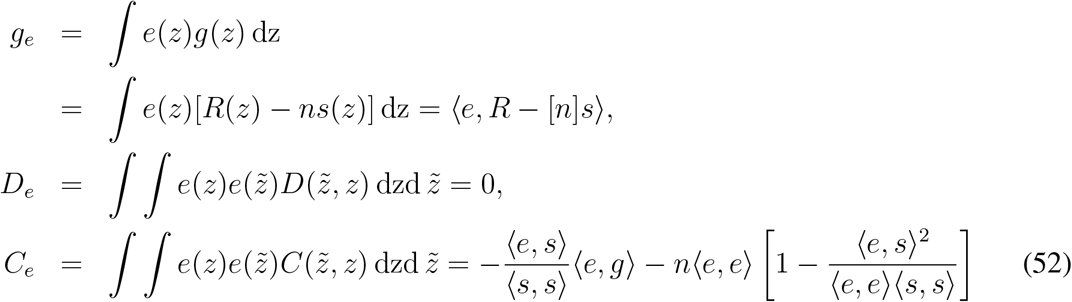

where [*n*] means that *n* is not a function of *z* but a scalar. When *g_e_* = 0 for any *e*(*z*), which corresponds to an evolutionary ideal free distribution (*R*(*z*) − *ns*(*z*) = 0 for all *z*), we see

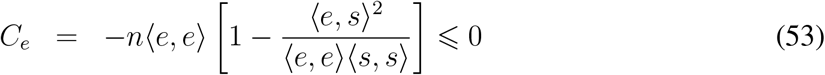

holds for any *e*(*z*). And *C_e_* = 0 holds only for *e*(*z*) ∝ *s*(*z*), which does not change the shape of *s*(*z*). Hence, an E-IFD (evolutionary ideal free distribution) is strongly convergence stable against mutations that change the shape of resource utilization distribution.

### 4.2 Evolution with constraints

#### 4.2.1 Constraints

We consider two constraints

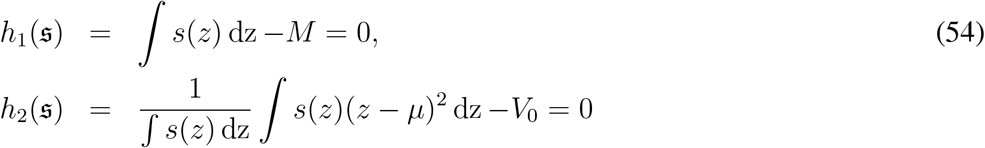

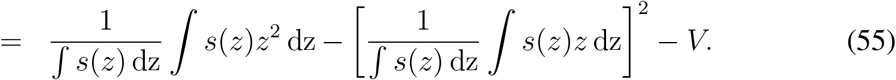

The 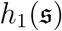 restricts the amount of the utilization distribution to *M*, and 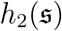 restricts the variance of the utilization pattern to *V*. Their first and second derivatives are given by

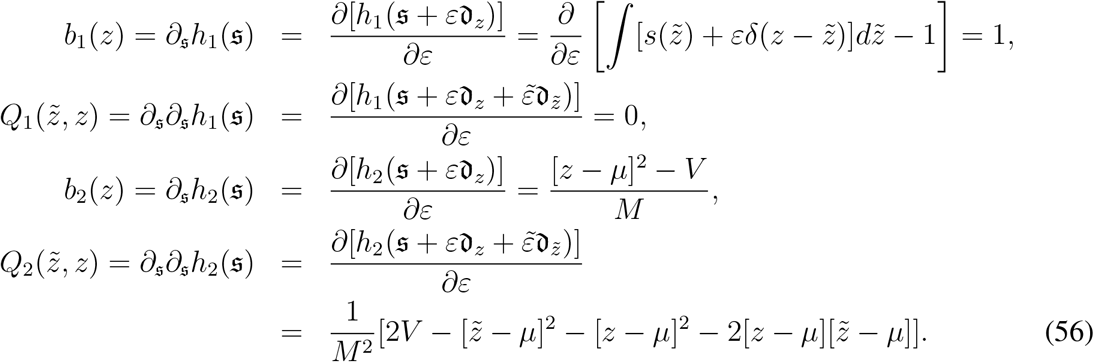

#### 4.2.2 Fitness gradient along constraint surface

The Lagrange fitness function is defined by

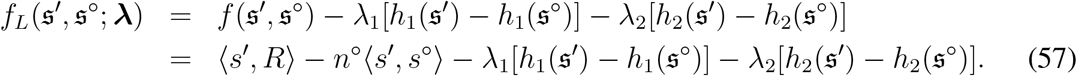

For a point 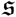 satisfying 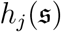 for all *j* = 1, 2, the λ = (λ_1_ λ_2_) is given by

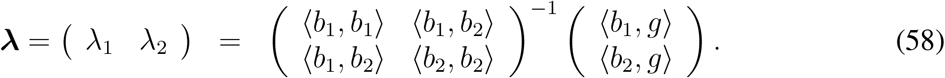

The fitness gradient at 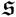 along the constraint surface is expressed as

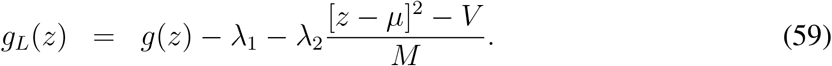

When 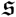 is evolutionarily singular, i.e., it has no fitness gradient along constraint surface, satisfying *g_L_*(*z*) = 0 for all *z*, we see

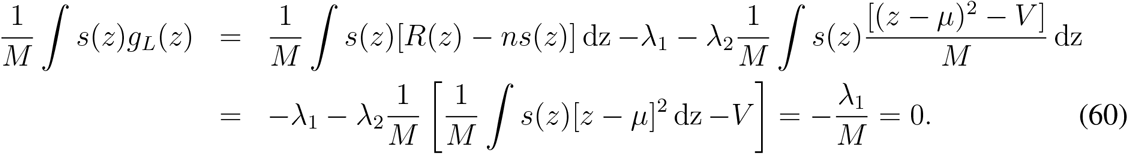

Moreover, we see

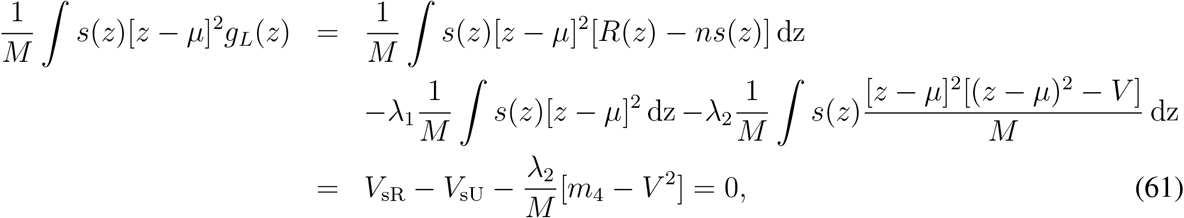

where

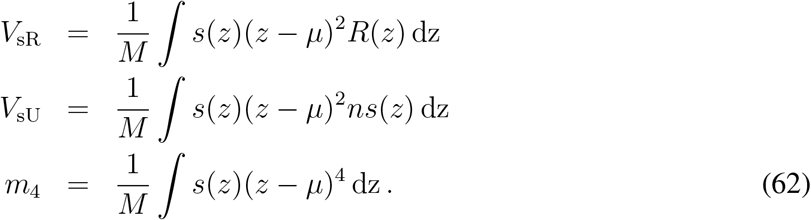

Thus, we get

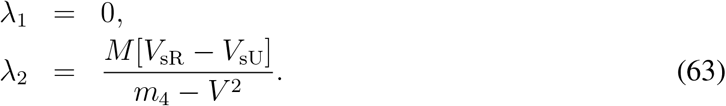

#### 4.2.3 Convergence stability and evolutionary stability along constraint surface

When 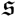 is evolutionarily singular, the *C_h_*(*e*) and *D_h_*(*e*) for *e*(*z*) satisfying ⟨*e, b*_1_⟩ = 0 and ⟨*e, b*_2_⟩ = 0 are given by

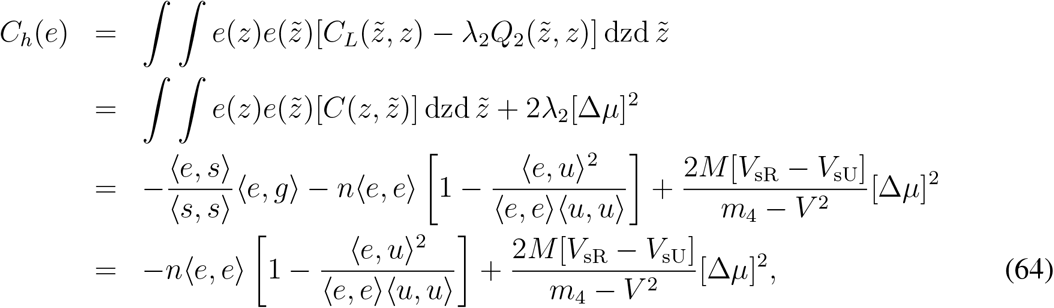

and

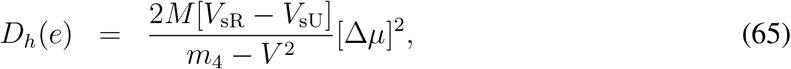

where

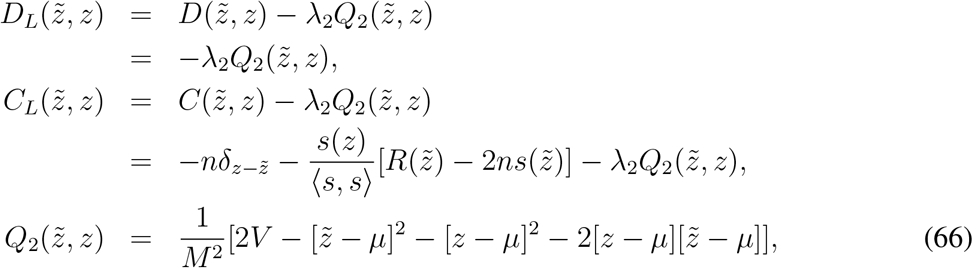

and

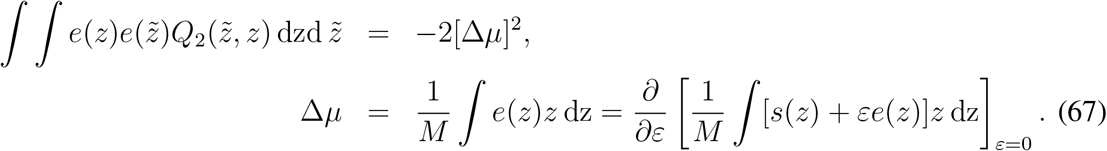

Therefore, if the visible resource variance *V*_sR_ is larger than visible total utilization variance *V*_sU_, then any *e*(*z*) that satisfies ⟨*e, b*_1_⟩ = 0, (*e, b*_2_) = 0 and Δ*μ* ≠ 0 satisfies *D_h_*(*e*) > 0, i.e., 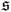 is evolutionarily unstable against the mutation *e*(*z*). In other words, 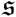 is evolutionarily unstable against all mutations that keeps the amount and variance of utilization distribution but changes its gravity center, as long as visible resource distribution is wider than the visible total utilization (by all existing individuals).

On the other hand, *V*_sR_ − *V*_sU_ *>* 0 decreases the convergence stability of 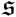. However, as far as |*V*_sR_ − *V*_sU_| is sufficiently small, i.e., the system is sufficiently close to an E-IFD, 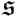 is strongly convergence stable. Hence, we can expect that an evolutionary singular point is almost always an CBP (candidate for evolutionary branching point) as far as the corresponding system state is sufficiently close to an E-IFD.

## 5 Discussion

This paper derives the conditions for candidate branching points (conditions for evolutionary singularity, strong convergence stability, and evolutionary instability), i.e., CBP conditions, for function-valued traits under simple equality constraints, by extending the CBP conditions for vector traits (Ito and Sasaki, 2016) for infinite-dimensional vector traits. As shown in the application example, the derived CBP conditions can provide considerably general insights, because function-valued traits are flexible. In principle, the CBP conditions are applicable for any constraint that has its first and second derivatives in the forms of *b*(*z*) and 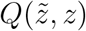, respectively. Investigation of what kinds of constraints have such property enables extension of the CBP conditions for more complex constraints (Dieckmann et al., 2006; Parvinen et al., 2006, 2013; Metz et al., 2016).

